# Convergence on reduced aggression through shared behavioral traits in multiple populations of *Astyanax mexicanus*

**DOI:** 10.1101/2022.05.02.490357

**Authors:** Roberto Rodriguez-Morales, Paola Gonzalez-Lerma, Anders Yuiska, Ji Heon Han, Yolanda Guerra, Lina Crisostomo, Alex C. Keene, Erik R. Duboue, Johanna Kowalko

## Abstract

Aggression is a complex behavior that is observed across the animal kingdom, and plays roles in resource acquisition, defense, and reproductive success. While there are many individual differences in propensity to be aggressive within and between populations, the mechanisms underlying differences in aggression between individuals in natural populations are not well understood. We addressed this using the Mexican tetra, *Astyanax mexicanus*, a powerful model organism to understand behavioral evolution. *A. mexicanus* exists in two forms: a river-dwelling surface form and multiple populations of a blind cave form. We characterized aggression in surface fish and cavefish in a resident/intruder assay through quantifying multiple behaviors occurring during social interactions. Surface fish, which are aggressive, display multiple social behaviors in this context, which we characterized into two types of behaviors: aggression-associated and escape-associated behaviors. The majority of these behaviors were reduced or lost in Pachón cavefish. Further, both aggression-associated and escape-associated behaviors were not dependent on the presence of light, and both surface fish and cavefish remained aggressive or non-aggressive, respectively, when opposed to fish from a different population. Additionally, we found that within populations, levels of stress response were not correlated with aggression- or escape-associated behaviors. Finally, when we compared aggression- and escape-associated behaviors across four cavefish populations, we found that both types of behaviors are reduced in three cave populations, while still present in one. Together, these results reveal that multiple cavefish populations have repeatedly evolved reduced aggression through shared behavioral components, while other cavefish have retained aggression.

**Summary Statement:** Comparison of aggression between surface fish and cavefish demonstrates that multiple complex behaviors compose aggression in surface fish and reveals heterogeneity in loss of aggression in cave populations.

## Introduction

Aggression is broadly defined as hostile behavior that creates harm or damage from one individual to another individual [1]–[3]. Motivation for aggressive behaviors can stem from multiple factors, including resource acquisition, establishment of hierarchies, survival and reproductive success [4]–[7]. Further, extreme levels of aggression can be detrimental for survival, suggesting aggression might be under stabilizing selection in some species [8] and highlighting the adaptive importance of regulating levels of aggression. Teleost fish are excellent models for studying aggression, as multiple species of fish display aggressive behaviors, including *Betta splendens* (Siamese fighting fish), multiple species of cichlids, sticklebacks, and the zebrafish *Danio rerio* [9]–[13]. While significant work in fishes has focused on the neural underpinnings of aggressive behaviors [14]–[16], the mechanisms contributing to evolution of aggressive behaviors are less well understood.

The Mexican tetra, *Astyanax mexicanus*, is a powerful emerging model for investigating the evolution of social behaviors [17]. *A. mexicanus* is a single species of fish consisting of river-dwelling, eyed and pigmented surface fish and at least 30 populations of blind, cavefish exhibiting reduced pigmentation or albinism [18], [19] Cavefish populations have evolved a number of behavioral differences relative to surface fish, including reduced sleep and schooling [20]–[22] increased vibration-attraction behavior (VAB) for prey detection [23], [24], and reduced aggression [25]–[28], providing a basis to investigate the ecological and genetic factors that underly the evolution of complex behavior. Cavefish and surface fish are interfertile, allowing for assessment of the genetic basis of behavioral evolution in this species through crosses and genetic mapping approaches [17]. Further, at least some cavefish populations have evolved independently of each other, providing the opportunity to examine whether cave-associated traits have evolved repeatedly [29], [30].

Teleost fish from other species exhibit a number of behaviors during aggressive encounters, including biting, striking, circling, following, escaping, freezing and avoidance [13], [27], [31]. While some of these behaviors have been reported in surface fish populations [32], previous work in *A. mexicanus* quantified aggression as a single metric, the number of attacks between pairs of fish [25], [26]. Thus, whether reduced aggression in cavefish is characterized by reduction in all or a subset of the multiple social behaviors that are exhibited during aggressive encounters in surface fish, and if these behaviors are under independent genetic control and thus evolve independently, is currently unknown.

To characterize the reductions in aggression that have evolved in cavefish, we quantify traits comprising aggressive-associated and escape-associated behaviors during social encounters of *A. mexicanus* surface fish and cavefish. Specifically, we asked: (1) What are the differences in the social behaviors that compose aggressive encounters in surface fish and cavefish, and are these behaviors dependent on sex, social context, or immediate environment? (2) Are the behaviors that occur during aggressive encounters repeatedly lost in multiple, independently evolved cave populations? Our findings position *A. mexicanus* as a powerful model for addressing how natural genetic variation contributes to a complex suite of aggressive behaviors, and how aggressive behaviors evolve.

## Materials and methods

### Fish Husbandry

All animal husbandry was performed according to methods previously described [33], [34]. All protocols were approved by the IACUC of Florida Atlantic University. Fish were raised at 23 ± 1°C. Adult *A. mexicanus* were housed in groups on a circulating filtration system in 18–37- liter tanks on a 14:10 hour light cycle that was constant through the animal’s lifetime. All fish used in this study were bred and raised in the laboratory. There were no statistical differences between surface fish from Río Choy and Texas lineages, and both populations were used in this study. Cavefish originated from the Pachon, Molino, Tinaja or Los Sabinos caves. All fish were 6 months – 1-year adults, which ranged from 3 to 6 cm in length.

### Resident-Intruder assay

All fish assayed for aggression were fed one hour before behavioral acclimation and assayed between zeitgeber time (ZT) ZT0-ZT6 Aggressive behaviors were quantified using a resident-intruder assay, which was previously shown to induce aggressive behavior in *A. mexicanus* and other vertebrates [31]. Pairs of resident and intruder fish from the same home tank were transferred to individual 2.5 L plastic tanks and acclimated for 18 hours in a dedicated behavioral room in which the light: dark cycle was maintained. All pairs of fish were sex- and size-matched. Following acclimation, the intruder fish was transferred to the tank of the resident fish and their interaction was recorded for 1 hour using a Microsoft Studio Webcam (#Q2F-00013). All recordings were performed from the front of the tank. For recordings in darkness, fish were acclimated and assayed in the dark. Infrared (IR) lights (850 nM) and cameras that could detect IR light were used during the resident-intruder assay. All resident-intruder recordings were acquired at 15 frames per second using VirtualDub2 (Version 1.10.5), an open-source video-capture and processing utility developed for Microsoft Windows (https://www.virtualdub.org/features.html).

### Novel tank assay

The novel tank assay, a well-established assay for assessing stress-like behaviors in fish, was performed on a subset of fish that were subsequently assayed for aggression in light versus dark conditions. All adult fish were of similar size (3-6cm). Stress assays were performed between Zeitgeber (ZT)6-ZT7 (ZT0=start of the light phase) as previously described [35], [36] with minor modifications. Groups of fish were transferred from their home tanks on the fish system into tanks in a dedicated behavioral room and allowed them to acclimate to the room for at least 1 hr. Next, each fish was transferred to a 500mL plastic holding tank for 10-minute acclimation, followed by gentle transfer into a 2.5 L tank containing 2 L of conditioned fish system water. Once transferred, fish were filmed in the light for 10-minutes using a Microsoft Studio Webcam (#Q2F- 00013). All stress recordings were acquired at 30 frames per second using VirtualDub2.After behavioral recording, the fish were housed individually in their respective tanks for acclimation in the resident-intruder assay.

### Manual Behavior Annotations

We annotated all staged-fights using the Behavioral Observation Research Interactive Software (BORIS) event-logging program [37]. For all annotations, we used the following ethogram based on previous behaviors observed in *A. mexicanus* and other fish species, and our own observations [13], [38], [39] (Table 1). Some behaviors were scored as single events in time (point events = biting, striking, circling) or continuous behavioral events (state events = following, escaping, freezing, avoidance). Individual fish behavior was scored throughout the video to distinguish between resident and intruder fish.

**Table 1.** Definitions for all aggression- and escape-associated behaviors scored in the resident/intruder assay.

| **Behavior** | **Description** |
| --- | --- |
| **Biting** | Focal fish physically makes contact with another fish with its mouth while performing an opening and closing motion with its mouth. |
| **Circling** | Both fish engage in a circular motion, typically with one head facing the tail of the other fish and vice versa. |
| **Following** | Focal fish follows the trajectory of another. |
| **Escaping** | Focal fish accelerates away from the other fish. |
| **Freezing** | Focal fish stops moving for greater than 5 seconds in any position within the tank. |
| **Avoidance** | Focal fish localizes in a corner of the tank for greater than 5 seconds. |
| **Striking** | Focal fish accelerates towards another fish ending in contact (but not necessarily biting). |

### Data Analysis

#### Manual Annotation in BORIS for Aggression

All data was exported from BORIS as activity plots and time budgets for quantification as text files (*txt) [37]. Time budgets represented the number of events during which a given behavior happened and the duration of such events, if applicable. For each time budget, the number of times each behavior happened was recorded for all behaviors, while the total duration (in seconds) was recorded only for the behaviors that had a time component (following, escaping, freezing and avoidance).

#### Automated Tracking for Stress

The center position of each fish was tracked using automated tracking with Ethovision software, and x-y displacement was calculated across all frames from the 10-minute recordings following previously published protocols using Ethovision XT13 (version 13.0, Noldus, Inc., Leesburg, VA) [40], [41]. To quantify bottom-dwelling for each fish, the arena was divided into three equal sections in Ethovision and the total duration of time spent in the bottom third of the arena was calculated. Ethovision accurately tracked the position of the fish using background subtraction.

Quantifications of all behaviors can be found in the supplementary materials.

### Statistical Analysis

We imported all data extracted from BORIS to GraphPad Prism 9. All data was tested for normality using Shapiro-Wilk test and parametric (t-tests for 2 group comparisons and One-Way-ANOVA for multiple group comparisons of a single variable) or non-parametric (Mann-Whitney for 2 group comparisons and Kruskal Wallis for multiple group comparisons) tests were used when appropriate. When analyzing more than one variable, such as the case when comparing the variation between light and dark conditions in surface fish versus cavefish populations, we used 2-Way-ANOVAs or Kruskal Wallis. Data was considered statistically significant if p < 0.05 (*), p < 0.01 (**), p < 0.001 (***), p < 0.0001 (****).

We used the Spearman’s rank-order correlation test to measure the association between all aggressive behaviors annotated with stress, and we calculated the rho (r_s_) for each correlation.

Outputs from statistical tests can be found in the supplementary materials.

## Results

### Aggression-associated Behaviors Are Observed in Surface Fish and Are Reduced in Pachón Cavefish

To characterize the behavioral repertoire that composes aggressive interactions in *A. mexicanus*, we performed a resident/intruder assay in surface fish (n = 10 pairs) and Pachón cavefish (n = 11) and annotated multiple behaviors displayed during aggressive interactions (Fig 1). Surface fish are highly aggressive, and pairs of fish exhibit a number of aggressive interactions throughout the course of the behavioral trial [25], [31], [42]. Surface fish display a number of behaviors observed in other fish species during aggressive interactions, including biting, striking, circling, and following (Fig 1A, B). While behaviors like fin fanning have been observed in other fish species, like in the Siamese fighting fish [43], we did not observe instances of these behaviors in *A. mexicanus* surface fish. In addition to these aggressive behaviors, fish exhibited a number of behaviors typically associated with subordinate/defeated status [13], [33], [44], including escaping, freezing and avoidance (Fig 1A, B). We quantified these behaviors in pairs surface fish and Pachón cavefish assayed in the resident/intruder assay. While Pachón cavefish exhibit some behaviors associated with both aggression and escape, most of these were significantly reduced in Pachón cavefish compared to surface fish. Pachón cavefish perform fewer aggression-associated behaviors compared to surface fish, including biting (p<0.05), striking (p<0.0001), and following (p<0.05) (Fig 1B-D). While surface fish also exhibit escape-associated behaviors, many of these were either reduced or absent in Pachón cavefish, including escaping (p<0.0001), freezing (p<0.001) and avoidance (p=0.01) (Fig 1F-H). Interestingly, both surface fish and Pachón cavefish exhibited circling behavior, and Pachón cavefish performed significantly more circling than surface fish (p<0.01) (Fig 1E), suggesting circling could be an aggression-associated behavior conserved in Pachón cavefish, or a social behavior serving another purpose in one or both populations of *A. mexicanus*. To assess whether sex played a role in the number of aggression- or escape associated behaviors observed in this assay, we performed a 2 way-ANOVA and found no significant effect of sex on aggression- or escape-associated behaviors, and no significant interaction between sex and population for any behavior, except for avoidance, where surface fish males were performing more avoidance than females (Fig S1, Supplementary Data sheets 1&3). Together, this suggests that reduced aggression in Pachón cavefish is characterized by reductions in a number of aggression-associated behaviors observed in surface fish.

**Figure 1.**
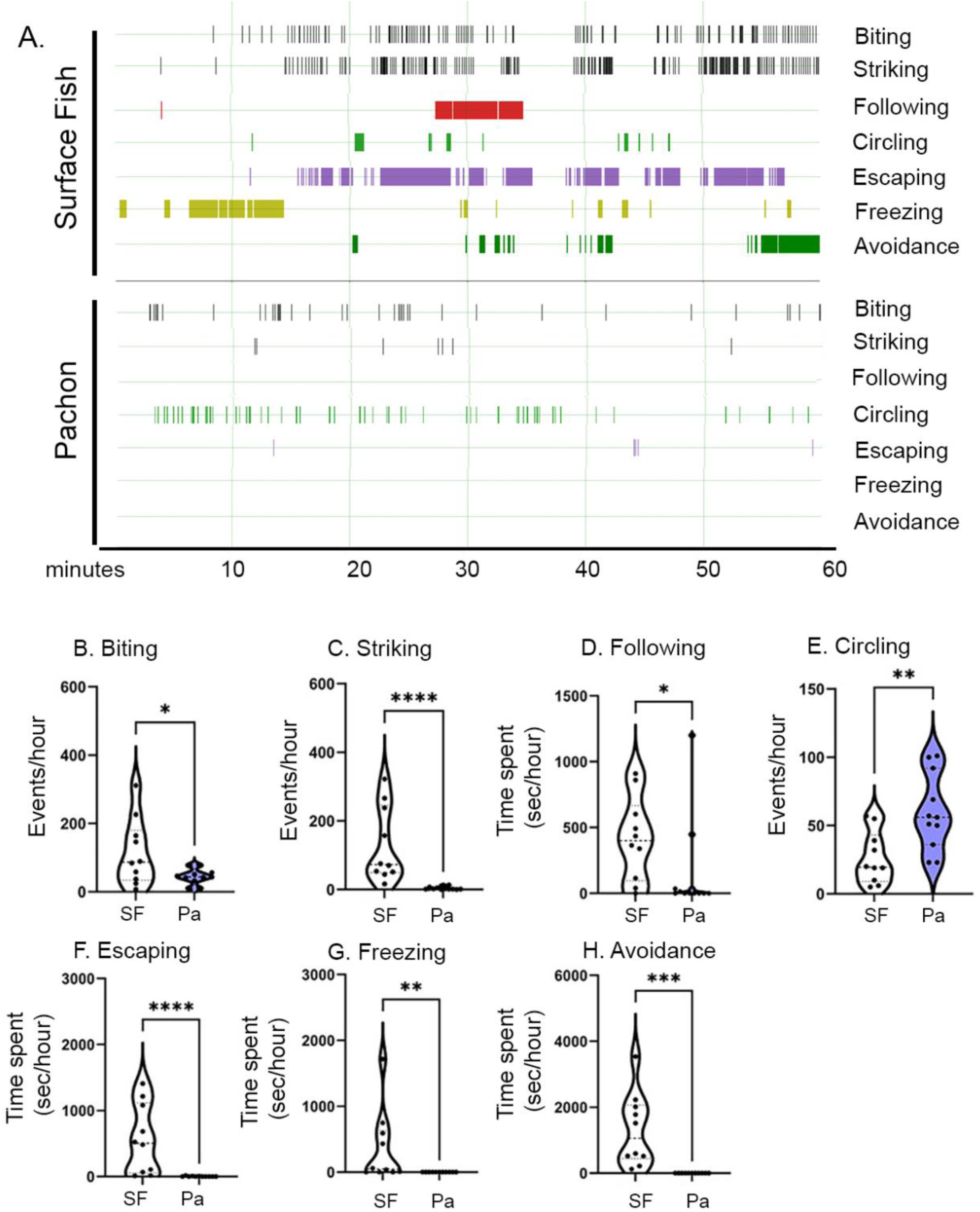
Quantification of social behaviors in the resident/intruder assay for surface fish and Pachón cavefish. (A) Representative ethograms for pairs of surface fish (top) and Pachón cavefish (bottom) during the resident/intruder 1-hour assay. Seven behaviors were annotated: biting, striking, following, circling, escaping, freezing, and avoidance (Table 1) over the 60 min assay period. Behaviors were quantified for each fish, but were pooled for both fish in each resident/intruder assay here (surface: n=10, Pachón: n=11). (B-H) Quantifications of behaviors annotated during the resident/intruder assay. All behaviors were scored for both individuals in the tank, and each data point represents either the number of behavioral events (biting (B), striking (C), circling (E)) or the time spent in a behavioral state (following (D), escaping (F), freezing (G), avoidance (H)) for one trial. Unpaired t-tests were calculated for biting (p<0.05), circling (p<0.01) and freezing (p<0.001). Mann-Whitney statistical tests were performed for striking (p<0.0001), following (p<0.05), escaping (p<0.0001) and avoidance (p<0.01). Significance: p < 0.05 (*), p < 0.01 (**), p < 0.001 (***), p < 0.0001 (****), not significant (ns).

Next, we asked whether aggression-associated behaviors were similar in quantity in both fish in each assay, or whether there was an asymmetry in how fish behaved, with one fish consistently acting as the aggressor and the other fish consistently escaping. As we tracked individual fish during our behavioral annotations, we examined whether there were quantitative differences in behaviors associated with resident/intruder status. We found no significant effects of resident/intruder status on any aggression- or escape-associated behaviors, or statistically significant interactions between resident/intruder status and population (Fig S2, Supplementary Data sheets 1&3). Next, we assessed whether within these assays, we could quantify behavioral symmetry between both contenders in each fight, regardless of resident/intruder status. To do so, we designated the fish in each pair that exhibited more strikes that aggressor, and the other fish the non-aggressor. When we compared aggression-associated and escape-associated behaviors for the aggressor versus non-aggressor in surface fish, we found there is a significant asymmetry in most aggression- and escape-associated behaviors in surface fish, with the aggressor performing significantly more biting, striking and following than the non-aggressor, and the non-aggressor performing significantly more escaping and avoidance than the aggressor (Fig S3). This pattern was not present in Pachón cavefish, consistent with the reduced aggression observed in fish from this population (Fig. S3). Together, these data suggest that, within pairs of surface fish, one fish is quantitatively more aggressive, and that this asymmetry is not observed in cavefish which have evolved reduced aggression.

### Aggressive Behaviors in Surface Fish and Pachón Cavefish Remain Constant under Light or Dark Conditions

Some social behaviors in surface fish, including schooling and shoaling, are reduced or absent when visual cues are not available [21]. To determine if this is the case for the aggression- and escape-associated behaviors quantified here, we performed resident/intruder assays under both light and dark conditions. Surface fish and Pachón cavefish exhibited similar behavior under light and dark conditions for the majority of the behaviors quantified (Fig 2). However, both surface fish and Pachón cavefish performed less circling in the dark relative to in the light (SF = 27.51 vs. 5.432, p<0.01, Pa = 41.19 vs. 19.11, p<0.01), suggesting that there is an effect of light dependency for at least one of these social behaviors (Fig 2F). Taken together, our data suggests that cavefish did not lose aggression simply due to the loss of the ability to receive visual cues to induce this behavior.

**Figure 2.**
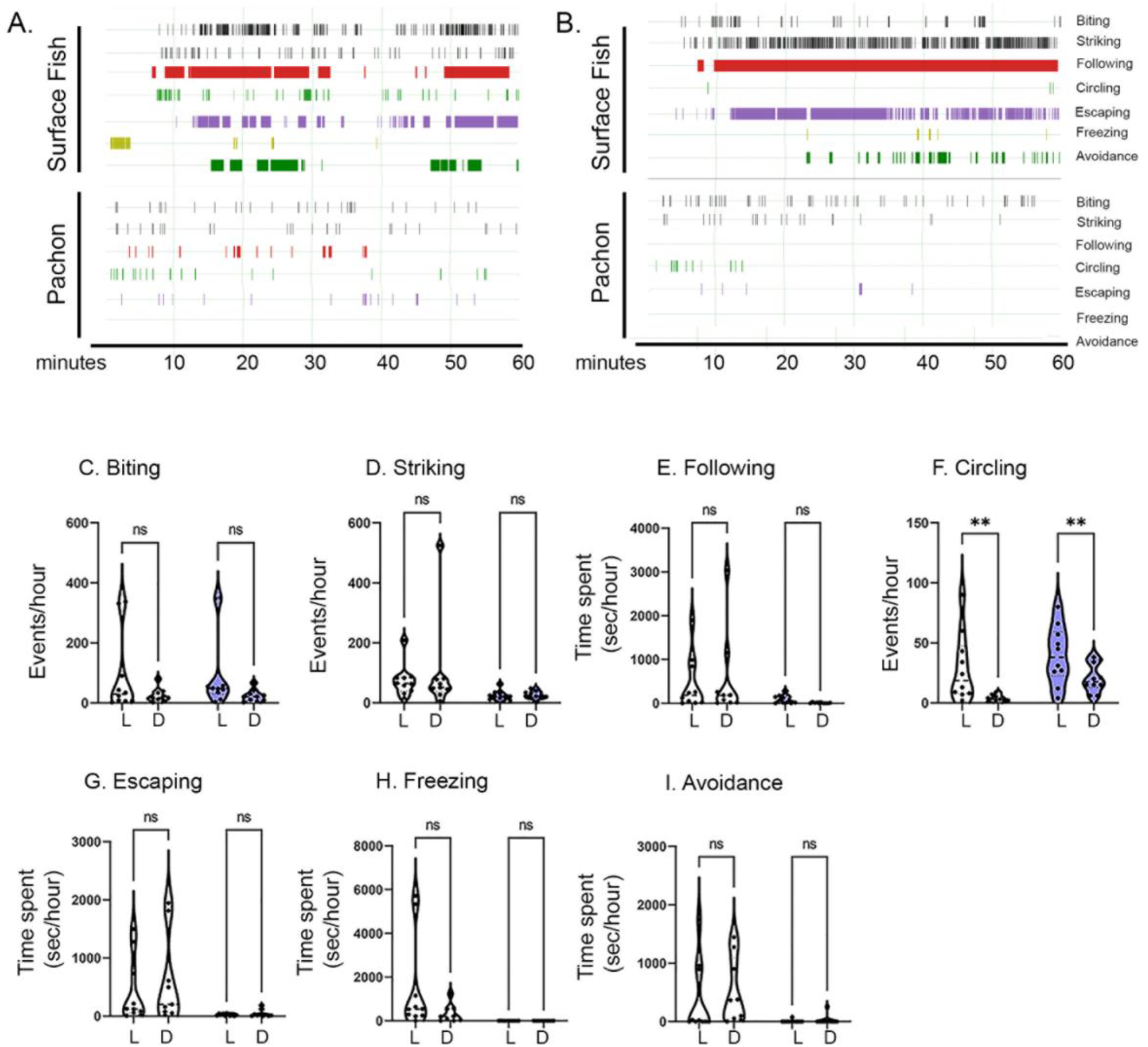
Social Behaviors in a Resident/Intruder Assay Under Light/Dark Conditions. (A-B) Representative merged resident/intruder activity plots for surface fish (top) and Pachón cavefish (bottom) in the light (A) or dark (B) during resident/intruder interactions. (C-I) Quantifications of behaviors annotated during each assay with light (L) versus dark (D) intra-population comparisons. 2-Way ANOVAs were performed for all behaviors in the light (surface fish, n = 10; Pachón cavefish, n = 10) and dark (surface fish, n = 9; Pachón cavefish, n = 10), followed by Tukey’s multiple comparison’s test for within surface or cave populations for light vs dark comparisons for biting (p=0.1169), striking (p=0.9446), circling (p=0.0063), following (p=0.9999), escaping (p=0.9020), freezing (p=0.4333), and avoidance (p=0.9337). Significance is reported only for comparisons within populations between light and dark: p < 0.05 (*), p < 0.01 (**), p < 0.001 (***), p < 0.0001 (****), not significant (ns).

### Surface fish Demonstrate Inter-population Aggression Towards Pachón Cavefish

It is possible that cavefish do not exhibit aggressive or escape-associated behaviors when they are interacting with other cavefish, but that these behaviors are inducible in the presence of another individual that exhibits them. To examine this possibility, we quantified behavior between inter-population pairs of fish in the resident/intruder assay under two conditions: (1) Surface fish-resident vs. Pachón-intruder (n = 8 pairs/each), and (2) Pachón-resident vs. Surface fish-intruder (n = 8 pairs/each). Surface fish exhibited aggression-associated behaviors when paired with a Pachón cavefish opponent (Fig. 3A-B), suggesting aggression is not associated with the identity the contender. These interactions induced one escape-associated behavior in Pachón cavefish, escaping (Fig. 3F). When surface fish were residents, they performed more striking and following than Pachón cavefish intruders, but this difference did not reach statistical significance (Fig. 3D, E). When Pachón cavefish were the residents, by contrast, most of the behavioral differences between resident and intruder observed were significant, with surface fish biting (p<0.01, Fig.3C), striking (p<0.01, Fig.3D) and following (p<0.001, Fig.3E) more that Pachón cavefish residents, and escaping less (p<0.01, Fig.3F). Taken together, this suggests that surface fish remain aggressive when opposed to cavefish, becoming even more aggressive when introduced as the intruders. Although Pachón cavefish do not become aggressive when opposed to a surface fish opponent, their interaction with surface fish induced escape-like responses, reminiscent of the profile of less-aggressive fish during surface fish contests (Fig S3).

**Figure 3.**
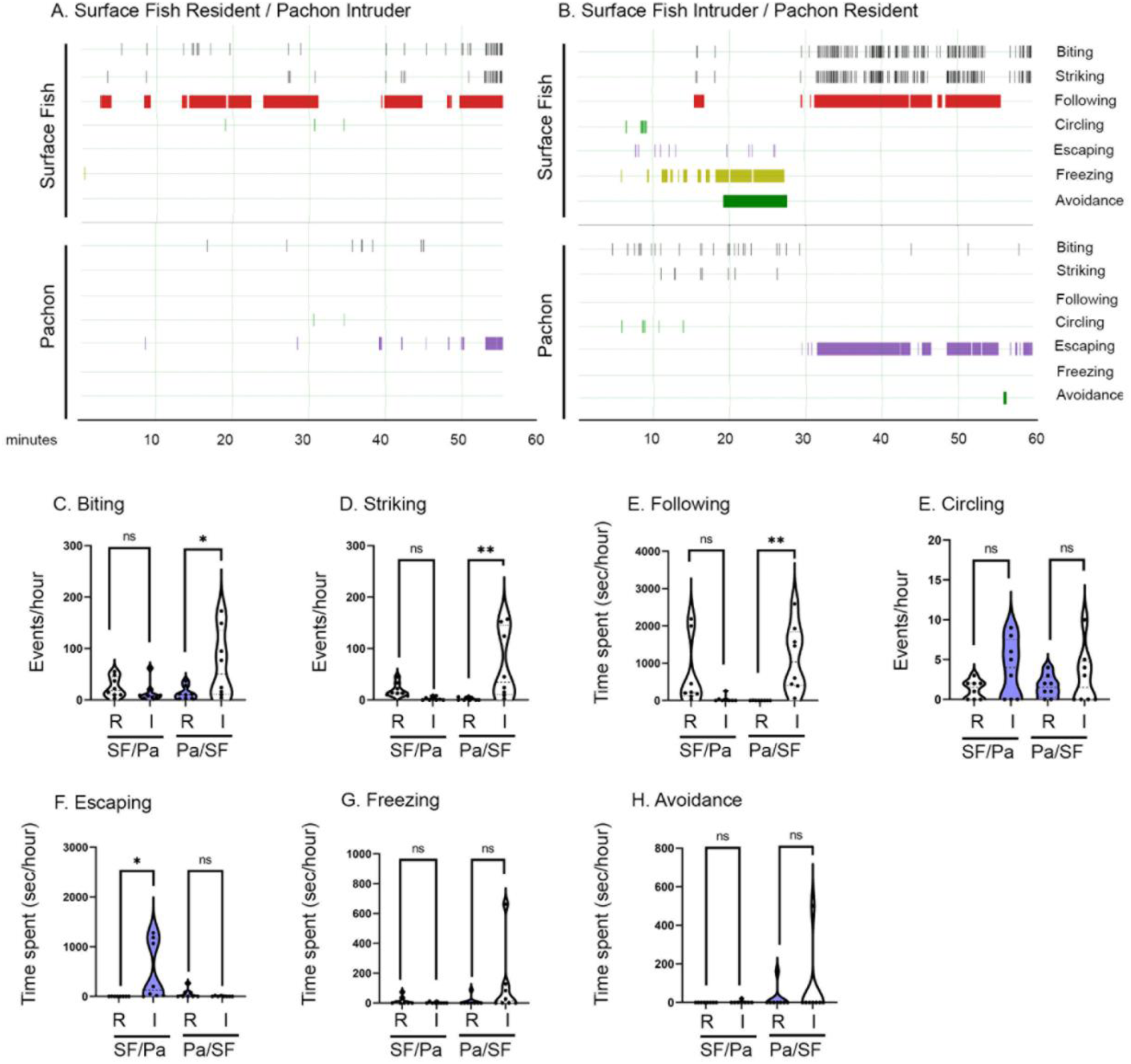
Resident/Intruder dynamics in surface fish versus Pachón cavefish fights. (A-B) Resident/intruder activity plots for surface fish-residents with Pachón-intruders (A) and Pachón- residents with surface fish-intruders (B) during staged fights. (C-I) Quantifications of behaviors annotated during staged fights with resident (R) and intruder (I) intra-population comparisons. 2-Way ANOVAs were performed for all behaviors, followed by Tukey’s multiple comparison’s test for resident versus intruder comparisons: When surface fish were residents: biting (p=0.9513), striking (p=0.7403), circling (p=0.9935), following (p=0.1689), escaping (p=0.9865), freezing (p>0.9999) and avoidance (p=0.9712). When Pachón cavefish were residents: biting (p=0.0293), striking (p=0.0060), circling (p=0.8589), following (p=0.0086), escaping (p=0.0245), freezing (p=0.2423) and avoidance (p=0.5680). Significance: p < 0.05 (*), p < 0.01 (**), p < 0.001 (***), p < 0.0001 (****), not significant (ns).

### Stress is Unrelated to Aggressive Displays in Surface Fish

Previous work suggested stress is an influencing factor on the onset of aggression [45]–[47]. To test this, we subjected surface fish and Pachón cavefish to an assay that has been used to quantify stress-related behaviors in multiple fish species [48]–[51], the novel tank assay [35], [52], prior to the resident/intruder assay acclimation for the comparisons of aggression in the light and the dark (Fig 2). As a proxy for stress, we measured the amount of time spent bottom-dwelling upon introduction to a novel environment, which was previously reported as a behavior exhibited when fish are stressed [53]. As reported previously, surface fish spend significantly more time at the bottom of the tank relative to cavefish (Fig S4). These observations confirmed previous findings that suggest surface fish are inherently more prone to stress than cavefish [41]. To examine whether some individuals within each of these populations exhibited more aggression-associated behaviors because they were more stressed, we compared the amount of time spent bottom-dwelling in the novel tank assay with the number of the aggression- or escape-associated behaviors we observed in fish in the light. We found no significant correlations between bottom dwelling and of the aggression- or escape-associated behaviors in either surface fish or in cavefish (Fig.4 and Fig.S5). Taken together, aggression appears to be unrelated to the stress profile within parental populations of fish, which suggests that differences in stress between cavefish and surface fish do not drive the differences in aggression observed.

**Figure 4.**
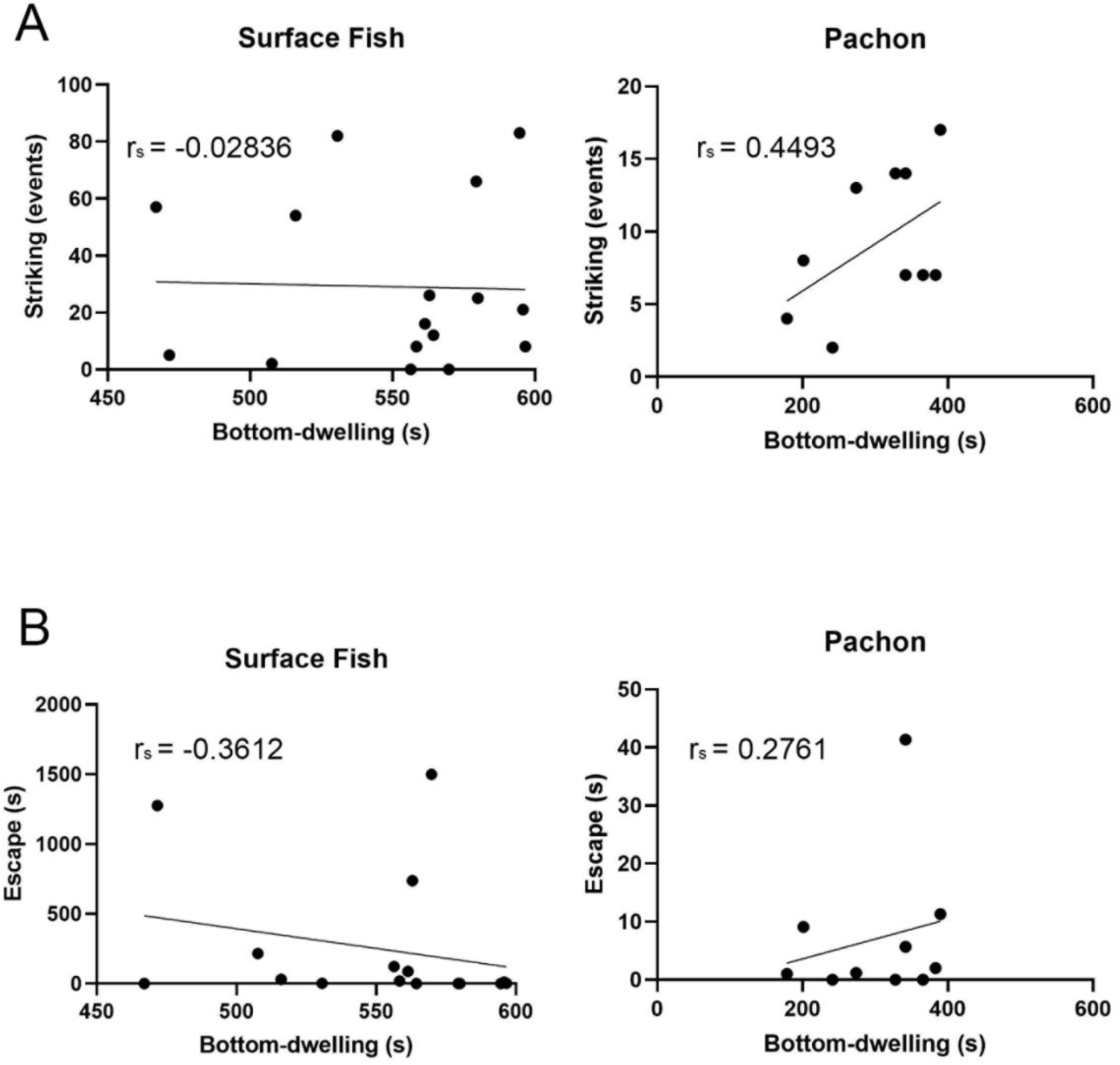
Correlation Between two social behaviors during a resident-intruder assay and bottom-dwelling. Correlations between number of strikes and time escaping during the resident/intruder assays and time spent in the bottom third of the tank in the novel tank assay were performed using Spearman’s rank correlation test for striking (A, surface, p = 0.9170<, Pachón, p = 0.1941) and escaping (B, surface, p = 0.1694, Pachón, p = 0.4416).

### Reduced Aggression is observed in cavefish from independently evolved cave populations

Populations of organisms that evolve under similar conditions often repeatedly evolve the same traits. *A. mexicanus* cavefish provide a powerful opportunity for studying repeated evolution, as multiple cavefish populations exist that have independently evolved a number of traits [17], [30], [54]. After assessing the repertoire of aggressive-like and escape-like behaviors which were present in surface fish and absent in Pachón cavefish, we asked if other cavefish populations have evolved reductions in aggression through modulation of the same aggression-associated behaviors. We quantified aggression in fish from four cavefish populations: Pachón (n = 5 pairs), Tinaja (n = 9 pairs) and Los Sabinos (n = 7 pairs) cavefish from the Sierra del Abra, and Molino cavefish (n = 12 pairs) from the Sierra de Guatemala. We found that Tinaja and Los Sabinos cavefish exhibited reduced or no instances of most aggression-associated and escape-associated behaviors, similar to the patterns found in Pachón cavefish (Fig.5). However, Molino cavefish exhibited a different set of behaviors compared to fish from these three cavefish populations. Specifically, Molino cavefish displayed more striking and more escaping than Pachón cavefish (Fig 5). Further, the increase in circling behavior we observed in Pachón cavefish relative to surface fish (Fig 1E), was not present in other cavefish populations (Fig 5A, E). These results suggest that reduced aggression has evolved in multiple, although not all, cavefish populations through reductions in multiple aggression-associated behaviors, and that some cave environments might favor the conservation of aggression-associated and escape-associated behaviors.

**Figure 5.**
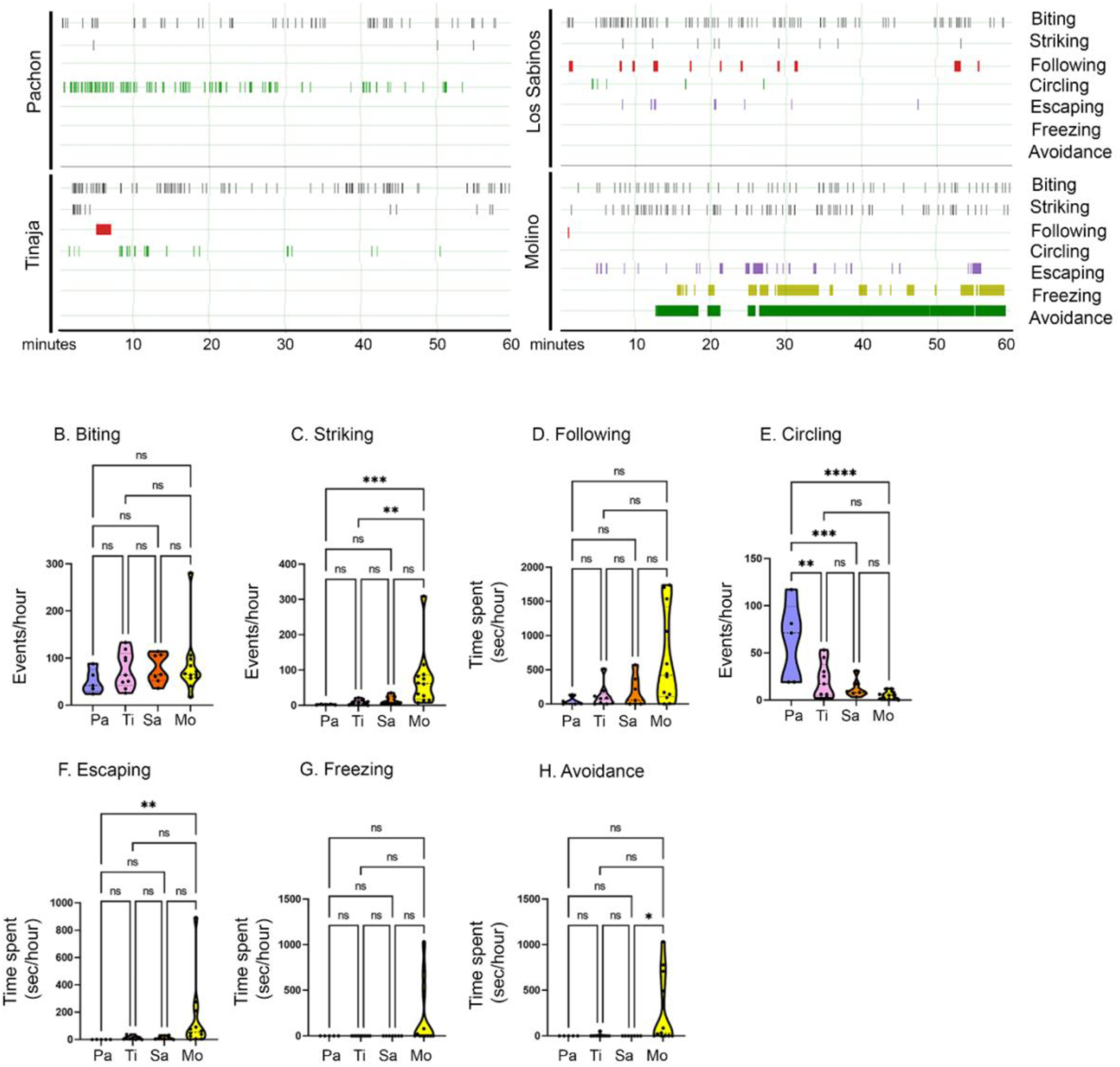
Social behaviors in a resident/intruder assay across multiple cave populations. (A) Representative resident/intruder activity plots for Pachón (top left), Tinaja (bottom left), Los Sabinos (top right) and Molino (bottom right) cavefish during the resident/intruder assay. The number of total behaviors for both the resident and the intruder were combined. All fish were sex and size matched, and sex was not used as a variable given the lack of effect of sex on seven behaviors in Pachón cavefish (Fig S1). (B-H) Quantifications of behaviors annotated during fights with comparisons across populations (Pachón = Pa, Tinaja = Ti, Los Sabinos = Sa, and Molino = Mo). One-way ANOVA followed by Tukey’s multiple comparisons test was performed for circling (Pachón-Molino, p<0.0001, Pachón-Tinaja, p<0.05, Pachón-Los Sabinos, p<0.001, Molino-Tinaja, p=0.2887, Molino-Los Sabinos, p=0.885, Tinaja-Los Sabinos, p=0.7954). Kruskal-Wallis with Dunn’s multiple comparisons test were performed for biting (Pachón-Molino, p=0.9213, Pachón-Tinaja, p>0.9999, Pachón-Los Sabinos, p=0.7564, Molino-Tinaja, p>0.9999, Molino-Los Sabinos, p>0.9999, Tinaja-Los Sabinos, p>0.9999), striking (Pachón-Molino, p<0.001, Pachón- Tinaja, p>0.9999, Pachón-Los Sabinos, p=0.2443, Molino-Tinaja, p=0.0057, Molino-Los Sabinos, p=0.3528, Tinaja-Los Sabinos, p>0.9999), escaping (Pachón-Molino, p<0.01, Pachón-Tinaja, p=0.409, Pachón-Los Sabinos, p=0.4466, Molino-Tinaja, p=0.206, Molino-Los Sabinos, p=0.341, Tinaja-Los Sabinos, p>0.9999), following (Pachón-Molino, p=0.0585, Pachón-Tinaja, p>0.9999, Pachón-Los Sabinos, p>0.9999, Molino- Tinaja, p=0.2307, Molino-Los Sabinos, p=0.3641, Tinaja-Los Sabinos, p>0.9999), freezing (Pachón-Molino, p=0.1938, Pachón-Tinaja, p>0.9999, Pachón-Los Sabinos, p>0.9999, Molino-Tinaja, p=0.0586, Molino-Los Sabinos, p=0.0995, Tinaja-Los Sabinos, p>0.9999), and avoidance (Pachón-Molino, p=0.0706, Pachón- Tinaja, p>0.9999, Pachón-Los Sabinos, p>0.9999, Molino-Tinaja, p=0.0702, Molino-Los Sabinos, p=0.0289, Tinaja-Los Sabinos, p>0.9999). Significance: p < 0.05 (*), p < 0.01 (**), p < 0.001 (***), p < 0.0001 (****), not significant (ns).

## Discussion

*Astyanax mexicanus* offers an opportunity to interrogate how complex behaviors, like aggression, evolve in closely related populations of fish. We took advantage of this and tested four blind cave populations (Pachón, Tinaja, Los Sabinos and Molino), which have evolved many traits independently, as well as sighted surface fish to probe for differences in aggression.

External factors from the environment can play a role in levels of aggression exhibited by individuals [55], [56]. Here, we examined whether morphological adaptations in cavefish, specifically loss of eyes and vision [26], were contributing to these differences in behavior. In other fish species, like the Coho Salmon, aggression is reduced in the dark, while in juvenile Atlantic Salmons serial reductions in light intensity decreased aggression [57], [58]. In *A. mexicanus*, there is some degree of controversy regarding aggression in the dark, as some studies report reduced aggression in surface fish in the dark [27], whereas others found that vision was dispensable for aggression in sighted surface fish [25], [26], and that surface fish raised following a lensectomy early in development are highly aggressive [26]. Our findings were in line with this latter work and further expanded this to demonstrate that multiple aggression-associated behaviors are observed under dark conditions. These differences in findings may be due to differences in the type of assay conditions [25], as well as the behaviors scored, and underscore the importance of quantifying a robust number of aggression-associated behaviors in this species.

Circling behavior has been associated with aggression in zebrafish [13], [59], sound-producing piranhas [60] and in *A. mexicanus* [31]. Our results were intriguing in the sense that Pachón cavefish perform fewer of all aggression- and escape-associated behaviors, except for circling. When compared with other cave populations, including Molino cavefish which exhibit aggression-associated behaviors, we found that increased circling is unique to Pachón cavefish. This behavior may not necessarily be aggression-associated, but instead serve a different purpose. For example, previous reports have found that Pachón cavefish perform stereotypic repetitive circling and that this behavior decreases under conditions that increase social interactions [61]. Thus, the circling behavior observed here in Pachón cavefish may not be used for social purposes. Alternatively, circling could be a social behavior in these fishes that is not associated with aggression.

Another question was if the presence of a fish from a different population triggered or suppressed aggression- and escape-associated behaviors. We tested this by adapting the resident/intruder assay for inter-population fights of surface fish and Pachón cavefish. These experiments led to two main findings: (1) surface fish were more aggressive than Pachón cavefish in intra-population assays, and (2) these interactions induced escape-associated behaviors in Pachón cavefish. While surface fish were were overall more aggressive than Pachón cavefish whether they were residents or intruders, these differences between surface fish and cavefish were larger when surface fish were intruders in the assay. This might be due to the existence of territoriality in surface fish, which has been proposed before [25], [27], [31], [62], and could result from surface fish intruders seeking to establish a territory. This finding was in contrast to our resident/intruder analysis of intra-population fights (surface versus surface, cave versus cave), where the fish’s designated role was not associated with whether the fish was ultimately the more aggressive or less aggressive fish in the assay. Hence, resident/intruder status does matter in this species, but this status appears to matter more when one fish is opposed to a fish from a different population. Our second finding here suggests while Pachón cavefish do not perform aggression- and escape-associated behaviors when interacting with other cavefish, they retain strategies to escape from an aggressive fish. Thus, aggression-associated behaviors may have been lost in Pachón cavefish while their escape-associated responses are still inducible. Evidence of cannibalism has been reported for *A. mexicanus* cavefish from the Micos cave [63]; thus, escape-like behaviors may be critical for survival in at least some cave environments.

Previous work introduced the notion that aggression and stress might be co-dependent behaviors in some fish species, or that at the very least, one of these behaviors could modulate the other [47], [64]. For example, in zebrafish, unpredictable chronic stress (UCS), as well as increases in stress-associated cortisol levels, increased aggression in male fish [64]. In *A. mexicanus*, stress-associated behaviors are reduced in the multiple populations of cavefish, including Pachón, Tinaja and Molino, relative to surface fish [36]. Further, intra-population differences in stress-levels, defined here as behavioral response to stress, were not correlated with levels of aggression- or escape-like behaviors in either cavefish or in surface fish, which suggests that, within *A. mexicanus* populations, individual differences in stress do not predict levels of aggression. Whether evolved differences in response to a stressful environment between populations is related to the evolution of reduced aggression in cavefish of this species remains to be determined.

Loss of aggression is observed in other animals that have evolved to live in cave environments, including other cavefish [65] and other cave species, like the whip spider *Phrynus longipes* [66]. To determine if and how aggression-associated behaviors have evolved across closely related cave populations, we examined whether repeated loss of aggression-associated behaviors has evolved in multiple cave populations of *A. mexicanus*. Studies in *A. mexicanus* using microsatellite and mitochondrial DNA suggest that cave populations are derived from at least two colonization events [67]–[70]. Surface fish of the “old stock” inhabited the Sierra de El Abra region and gave rise to the “old stock” of cavefish, including Pachón, Los Sabinos, Tinaja and others [71]. A different wave of surface fish gave rise to the present surface fish in the region and the “new stock” of cavefish, including Molino and Escondido [71]. Genetic studies suggest that many traits have evolved repeatedly in these different cave populations, whether they derive from these different colonization events, or even between cave populations from the El Abra caves. These traits including genetically encoded morphological traits such as the size of the eye primordia [72], [73], and behavioral traits, including foraging behaviors [74].

Recent work suggested Molino cavefish were not aggressive, and behaved similarly to Pachón cavefish, differing only in their patterns of attacks [25]. We observed that Molino fish show increases in at least one aggression-associated behavior relative to Pachón cavefish, which is in line with a previous study that found that Molino cavefish are aggressive [26]. This result could mean the ecological environment of the Molino cave favors the conservation of aggression- and escape-associated behaviors. Ultimately, these findings pose several new questions: (1) is “cavefish aggression” unique to Molino, or have other *A. mexicanus* cavefish conserved these behaviors? (2) Are these conserved aggressive behaviors specific to the cavefish derived from this colonization, or are other, currently untested cavefish populations from the Sierra de El Abra aggressive? (3) Do the same genes underlie reduced aggression in the Pachón, Tinaja and Los Sabinos populations? Sampling fish from more caves will provide answers to some of these questions. Ultimately, identifying and functionally interrogating the genes that are contributing to the loss of aggression in *A. mexicanus* will provide additional insight into the genetic factors contributing to natural variation in aggression in this species. Methods such as QTL analysis and functional interrogation of candidate genes using CRISPR-Cas9 that are available in this species could be used in the future to answer these questions [75]–[78]. Thus, this work provides a platform for investigating the extent to which heredity and/or environmental pressures inform the evolution of aggression across closely related populations in a same species.

